# Cannabidiol dose modulates behavioral response to acute and repeated administration of Δ9-Tetrahydrocannabinol by strain and sex

**DOI:** 10.64898/2026.03.05.709920

**Authors:** Sheena Saxena, Elizabeth A. Duecker, Lukmon M. Raji, Christine E. Watkins, Sufiya Khanam, Byron C. Jones, Bob M. Moore, Megan K. Mulligan

**Author notes:** Shared first author (Sheena Saxena, Elizabeth A. Duecker).

## Abstract

Cannabis contains many bioactive compounds, including Δ9-Tetrahydrocannabinol (THC) and cannabidiol (CBD), which influence behavior through complex pharmacological interactions with endogenous targets. This study examines whether CBD influences THC-induced changes in motor activity, hypothermia, and antinociception traits across different THC:CBD ratios, sexes, and genetic backgrounds. Traits were measured in C57BL/6J (B6) and DBA/2J (D2) mice of both sexes following baseline intraperitoneal (*i.p.*) injection of vehicle (VEH) and two consecutive daily doses of VEH or THC (10 mg/kg) alone or in combination with 0.56, 5, or 10 mg/kg CBD (THC:0.56CBD, THC:5CBD, or THC:10CBD, respectively). Motor activity and hypothermia were quantified daily from 0 to 120 min following injection and antinociception was measured daily at 60 min. We found that CBD alters THC-induced changes in motor activity and hypothermia as a function of day, dose, time, sex, and strain. In D2 females, CBD dose-dependently attenuated the hypolocomotor effects of THC immediately following acute injection and enhanced these effects later at 75 min. Following repeated exposure, CBD dose-dependently enhanced THC-induced hypolocomotion in B6 females at 75 min and in D2 males at 30 min while attenuating THC-induced hypolocomotion in D2 females immediately following injection. In D2 females, CBD dose-dependently attenuated THC-induced hypothermia at 15 min and enhanced hypothermia relative to THC at 30 min in D2 males following acute injection. After repeated exposure, CBD dose-dependently enhanced THC-induced hypothermia in B6 females at 15 min and in D2 males from 30 to 120 mins, while attenuating hypothermia in D2 females at 30 min. No significant effects of CBD on antinociception were observed. Our results indicate that CBD can modulate some THC-induced traits acutely and after repeated exposure. Regulation of THC-induced behavioral responses is dependent on CBD dose, genetic background, and sex. A candidate gene search using brain gene expression in recombinant inbred mice revealed greater genetic variation in ion channel genes relative to key metabolic genes, suggesting an underlying pharmacodynamic mechanism. Future research and validation of molecular mechanisms underlying these differences is expected to enhance our understanding of potential health risks and clinical relevance of cannabis and cannabinoid compounds containing THC and CBD.

## INTRODUCTION

The legalization of medicinal and recreational cannabis has resulted in the generation of various cannabis cultivars typically characterized by their unique chemical profiles (*e.g.*, cannabis chemotypes or chemovars). The foundational system for classifying cannabis chemotypes relies heavily on the ratio of just two of the hundreds of native phytocannabinoids—THC and CBD—although classification based on other constituents (*e.g.*, terpenes) has also been applied (1). The THC:CBD ratio has been associated with differences in psychoactive and/or alleged beneficial effects (2) of cannabis cultivars. For example, high-THC chemovars are often associated with higher intoxication and psychoactive effects compared to high-CBD chemovars, while intermediate effects are ascribed to more balanced levels of THC and CBD. Behavioral responses to a select strain of cannabis can be highly variable among individuals (3, 4) and may reflect the relative levels of THC and CBD, in addition to the abundance of other phytochemicals. Even though the phytochemistry of cannabis is complex (5, 6) there is compelling evidence from human and rodent studies that the THC:CBD ratio is an important determinant of behavioral response.

Numerous clinical and preclinical studies have compared the relative effects of THC, CBD, and combinations of both mixtures on behavioral and physiological outcomes (7-42). Relative to all possible dose combinations, ratios close to 1:1 are more frequently evaluated in comparison to either high or low THC:CBD cannabis chemovars or THC alone (9, 16, 20, 22, 23, 26, 27, 30, 35, 37-39, 41). In some studies balanced levels of THC and CBD were associated with attenuation of THC-induced effects, including psychotomimetic effects (16), anxiety (8, 13, 17, 39) memory deficits (8, 13, 17, 38, 39) and conditioned aversive responses (9). Equal ratios of each cannabinoid were also associated with the moderation of THC-induced effects on brain activity and blood flow (24). Varying the doses of THC and CBD beyond a simple 1:1 ratio in rodents was found to produce complicated dose-dependent effects on behaviors such as social interaction (11) and antinociception (12, 14). In human studies, coadministration of CBD by inhalation, intravenous, or oral routes of administration dose dependently attenuated THC-induced symptoms of anxiety or paranoia (8, 10, 34, 39) psychosis (10, 16) and the appetitive and reinforcing effects (43) but did not always diminish feelings of euphoria, intoxication or feeling high (16, 40). It should be noted that dose, route of administration, species, sex, and genetic background are highly variable among studies and can obfuscate direct comparisons. Despite the limitations, these studies demonstrate that CBD levels are important determinants of the pharmacological effects of THC and may underlie individual differences in response. Although there is evidence that CBD levels can influences behavioral outcomes attributed to THC, the precise underlying molecular mechanisms and the possible contribution of individual differences (*e.g*., age, sex, genetic background) remains unclear.

The complex pharmacology of THC and CBD provides many possible mechanisms through which variation in the relative levels of each might contribute to disparate effects on organismal response. Both phytocannabinoids are bioactive substances that can interact with the endogenous endocannabinoid system (ECS) as well as other endogenous targets to produce diverse effects through engagement of unique targets or differential agonism/antagonism of shared receptor-mediated signaling pathways (44). The primary ECS targets of THC are the G-protein coupled cannabinoid receptors 1 (CB1) and 2 (CB2) where it acts as a partial agonist. The psychotropic effects of THC in humans (*e.g.*, feeling high) and the behavioral effects in rodents (*e.g.*, analgesia, hypothermia, and changes in motor activity) have been attributed to CB1 activation in the central nervous system (45). Moreover, genetic or pharmacological deletion of CB1 abolishes these behavioral responses in rodents (45-49). However, Wang and colleagues recently reported that deletion of CB2 in rodents reduced the analgesic properties of THC without impacting motor or hypothermic effects, suggesting an additional role for CB2 in analgesia (50).

Unlike THC, CBD is non-psychotropic, and does not bind orthosteric sites in CB1 and CB2 (51). Paradoxically, CBD can block the effect of CB1 and CB2 activators (*e.g*., C9-55940 and WIN55212) at mouse CB1 and human CB2 (52, 53) suggesting that CBD may modulate both receptors through allosteric regulation (54), albeit at potentially higher levels than may occur *in vivo*. In addition, CBD has many unique endogenous targets and has been shown to activate serotonin receptors, modulate TRPV1 receptors, and inhibit the ECS catabolic enzyme FAAH (55, 56), among others. Indeed, several studies have suggested that CBD modulation of THC-induced behavioral outcomes may be best explained by a pharmacodynamic mechanism involving CB1, 5-HT1A, and/or TRPV1 (16, 27, 28, 30, 39, 57-61). However, human and rodent studies have also demonstrated inhibition of THC metabolism by CBD resulting in increased plasma THC levels and a reduction in active metabolites (7, 12-14, 36), suggesting possible involvement of pharmacokinetic mechanisms as well.

Identification of individual differences in the pharmacodynamic and/or pharmacokinetic mechanism(s) underlying the variable effects of THC:CBD ratios on behavior is important for understanding the physiological effects of cannabis as a whole. Elucidation of these mechanisms requires quantification of the impact of different CBD levels on THC-induced behavioral outcomes across different sexes and genetic backgrounds. To this end, our group previously examined the effects of murine genetic background and sex on characteristic behavioral responses (*i.e*., hypothermia, hypolocomotion, and antinociception) to high dose THC (10 mg/kg, *i.p.*) following acute or repeated exposure. We demonstrated that highly genetically divergent B6 and D2 inbred mouse strains exhibited striking sex and strain differences in the response to THC (62). Specifically, we identified strain differences in the level of THC-induced motor suppression (B6 > D2) and antinociception (D2 > B6) and sex differences in the level of THC-induced hypothermia (females > males) following acute exposure. We also demonstrated rapid tolerance to the THC-induced decreased in motor activity and body temperature, even after two consecutive daily exposures (62). Moreover, the D2 strain demonstrated greater rapid tolerance to THC-induced hypothermic and antinociceptive effects (62). In other work, we demonstrated that strain differences in THC-induced hypolocomotion and hypothermia following acute exposure are highly heritable (63), indicating the presence of segregating genetic variants between B6 and D2 mice in downstream signaling pathways that are capable of producing distinct adaptive responses to THC. In this paper we apply a THC and CBD coadministration paradigm to our previously described approach to determine whether CBD dose can modulate foundational behavioral responses (*i.e*., hypothermia, hypolocomotion, and antinociception) to THC. Specifically, we test whether different doses of CBD can enhance or attenuate THC-induced changes in these responses following acute or repeated exposure to THC. We also evaluate whether genetic background and sex influence the ability of CBD to modulate THC-induced behavioral responses and nominate potential candidate genes using a genetics approach.

## MATERIALS and METHODS

### Animals

C57BL/6J (B6, JAX #000664) and DBA/2J (D2, JAX #000671) mice were purchased from The Jackson Laboratory and allowed to acclimate for at least two weeks before experimental procedures. Mice were between 62–170 days old at the time of testing (median: 116 days; mean ± SD: 114 ± 27 days). Animals were maintained on a 12-hour light: dark cycle (lights on at 0600 h) with *ad libitum* access to standard chow diet (Inotiv 7912 fixed formula, irradiated diet) and water. All procedures were performed during the light cycle (0700–1600 h) in accordance with the University of Tennessee Health Science Center Institutional Animal Care and Use Committee (IACUC).

### Formulations and Drug Administration

THC and CBD were obtained from the National Institutes of Health (NIH) National Institute on Drug Abuse (NIDA) Drug Supply Program. THC and THC + CBD formulations were prepared in an ethanol:cremophor:saline (5:5:90) vehicle (VEH) and filter sterilized. Formulations were stored in septum-sealed vials at 4°C in the dark and used within one week. THC was dosed at 10 mg/kg, and THC+CBD combinations were formulated as follows. THC:0.56CBD was formulated as 10 mg/kg THC + 0.56 mg/kg CBD, THC:5CBD was formulated as 10 mg/kg THC + 5 mg/kg CBD, and THC:10CBD was formulated as 10 mg/kg THC + 10 mg/kg CBD. VEH and all formulations were administered at the same volume (100 µL per 30 g body weight). Stability testing indicated that THC retained 95.2% of its initial concentration after eight days of storage at 4°C (62).

### Experimental Design

Mice were randomly assigned to a treatment group. All mice (*n* = 177) were tested in 8 cohorts (**Supplemental Table 1)**. Cohort testing dates occurred in October of 2018 for cohort 1, November of 2018 for cohort 2, January of 2019 for cohort 3, February of 2019 for cohorts 4 and 5, March of 2019 for cohort 6, April of 2019 for cohort 7, and February of 2025 for cohort 8. Initial pilot studies were conducted with THC:0.56CBD in cohorts 1 through 3 to test whether the THC:CBD ratio commonly associated with cannabis-derived compounds produced large effect differences in response relative to THC alone. After finding minimal differences, we then expanded the study to include increasing amounts of CBD (*e.g*., THC:5CBD and THC:10CBD) in later cohorts.

All experiments were conducted during the light phase of the circadian cycle, typically between 9:00 and 17:00 h. Mice were implanted with subcutaneous RFID transponders (TP-500 Temperature Programmable Transponder 9mm, Avidity Science) at least four days prior to handling. All mice were housed alone for three days prior to handling or testing. To acclimate mice to handling and reduce aggression and escape behavior, mice were transported to the test room for at least three consecutive days prior to the study. During each day mice were allowed to acclimate for 30 min and then scanned for an RFID, transported via plastic container to a scale, and then transferred by hand to a plastic restraining cylinder used for the antinociception assay. Following handling, mice were returned to the home cage and transported back to the mouse colony. To habituate mice to the phenotyping pipeline, on the day before testing, mice were transported to the test room and allowed to acclimate for 30 min before placing in the open field for 10 minutes. Thirty minutes later, tail flick latency was assessed, and the mice were then returned to the colony. During the habituation phase, measurements were recorded but not used for statistical analysis.

The phenotyping pipeline (**Figure 1**) included measurements of body temperature, motor activity, and antinociception collected over three consecutive days at discreet time points following acute intraperitoneal (*i.p.*) injections of VEH, THC or THC+CBD formulations. Body temperature and subject ID were monitored using RFID transponders. Motor activity was measured over 10 min in a 40cm x 40cm x 40cm plexiglass open field and time spent mobile was tracked in the open field using AnyMaze software (Stoelting). Immobility was scored as no movement in the open field for 3 or more seconds. Antinociception was quantified as the latency to tail flick in response to a thermal stimulus in a 52°C water bath. Each day, temperature was recorded at 0-, 15-, 30-, 60-, and 120-min post-injection (VEH, THC, and THC + CBD), motor activity was recorded at 0-, 30-, and 75-min post-injection, and tail flick latency was recorded 60 min post-injection. Baseline measurements were recorded for each subject following a 30 min acclimation period to the testing room by giving all groups an injection (*i.p.)* of VEH and then recording temperature, time mobile in the open field, and tail flick latency at the previously described time intervals. On the second and third days, response to treatment was measured by recording temperature, motor activity, and antinociception after administering VEH, THC, or the three different combinations of THC + CBD.

**Figure 1.**
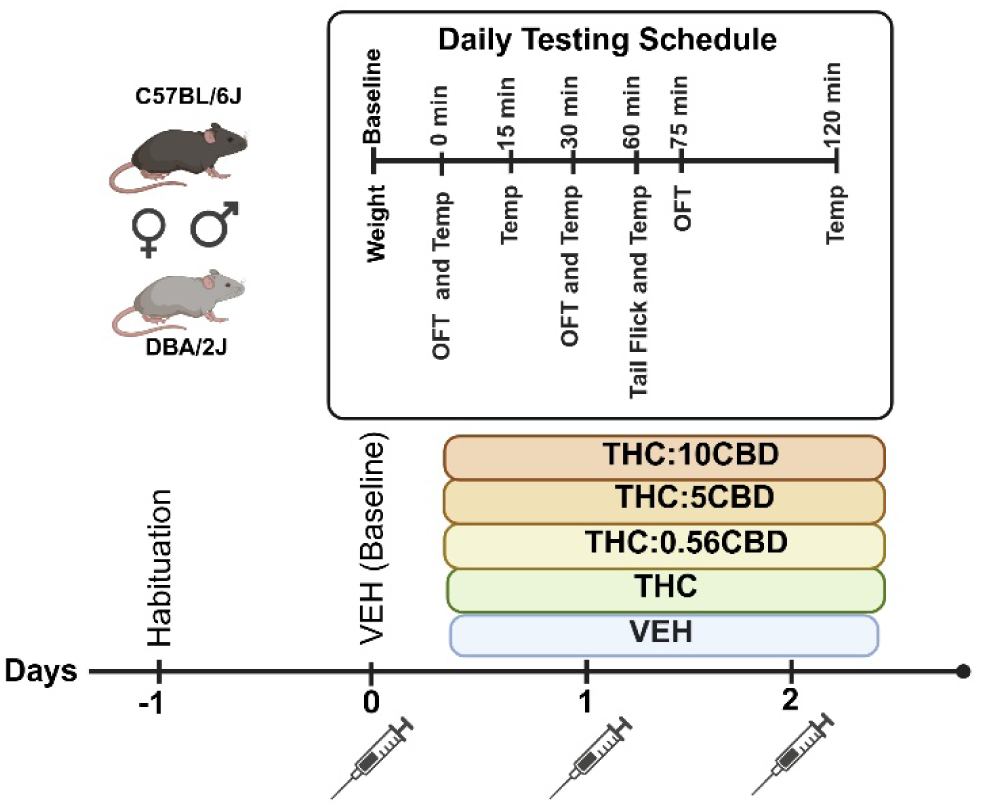
Overview of the experiment. Mice were assigned to one of five treatment groups and measurements of body temperature (temp), motor activity in the open field test (OFT), and antinociception (tail flick) were collected over three consecutive days at discreet time points following acute intraperitoneal (i.p.) injections of VEH, THC or THC+CBD formulations. All mice were habituated to the experimental pipeline on day -1, but no injections were given and the data was not used in the analysis.

### Statistical Analysis

#### Acute and repeated sensitivity calculations

To identify whether there was an effect of THC:CBD ratio on trait response to THC within strain, sex, time point and day, we calculated acute response following one or two consecutive daily exposures to VEH, THC, THC:0.56CBD, THC:5CBD, or THC:10CBD. Acute sensitivity was calculated as the difference between each trait response at baseline (*i.e.*, after VEH injection on day 1) and the first exposure to treatment (*i.e.*, VEH, THC, or THC + CBD mixtures) for each subject and time interval. Repeated sensitivity was calculated as the difference between each trait response at baseline and the second exposure to treatment for each subject and time interval. Previously, we found that B6 and D2 mice of both sexes show rapid tolerance to THC (10 mg/kg, *i.p.*) in the form of diminished THC-induced hypolocomotion, hypothermia, and antinociception following two consecutive daily exposures (62). *Treatment effects.* We first tested whether there was any significant difference in treatment means for initial and repeated sensitivity within each group (*e.g.*, sex, strain, and time interval) using one-way ANOVA with 5 levels of treatment (*e.g.*, VEH, THC, THC:0.56CBD, THC:5CBD, THC:10CBD). If there was evidence in support of a significant (*p* < 0.05) effect of treatment, we then used Dunnett’s post-hoc test to determine which treatments differed from either VEH or THC for each group. To do this we performed two different contrasts for all groups using Dunnett’s test; first we compared each treatment mean to VEH and then we compared the mean of each of the three THC + CBD groups to THC alone. These contrasts allowed us to distinguish the impact of THC and CBD dose on trait variation. For graphing purposes, we averaged the two difference scores for acute and repeated sensitivity within strain, sex, treatment, and timepoint and presented error as the standard error of the mean.

### Candidate Gene Search in GeneNetwork

Over 75 targets have been proposed to mediate the biological activity of CBD (64). Many of the reported targets are derived from *in vitro* activity analysis with EC_50_, IC_50_ or K_i_ values ranging from nanomolar to high micromolar. However, *in vivo* target exposure to CBD may differ significantly relative to *in vitro* data thus it is critical to recognize the effective concentration (C_max_) of a drug in the target tissue. Pharmacokinetic studies of CBD in rodent models have yielded C_max_ values ranging from 0.01 to 28.5 µM, dependent on route of administration and rodent species (7, 65-68). Bih and colleagues addressed the potential targets of CBD in the CNS utilizing the data reported by Deiana *et al. (65)* and set the upper limit of *in vitro* activity (EC_50_, IC_50_ or K_i_) to a range less than 10–20 µM (69). In the study cited by Bih, the dose of CBD was 120 mg/kg which in mice yielded a brain C_max_ of 1.3 mg/g p.o. (4.3 µM, using a *r_brain_* = 1.04 g/ml, (70)) and 6.9 mg/g i.p. (23 µM). At lower doses Ferreira *et.al.* (67) report a C_max_ of 0.33 µM with a dose of 10 mg/kg (i.p.) and 12.3 µM following *i.p.* administration of 50 mg/kg. Since our studies used CBD concentrations between 0.1-10 mg/kg, we used an upper limit of 10 µM to select targets of CBD proposed by Bih (69) a more recent 2025 review by Manzoni *et al.* (71). We performed additional literature searches to further refine the list of targets based on recent scientific literature. We ultimately selected seven targets proposed by Bih et al., (69) with an additional 21 from Manzoni *et al.* (71), and one additional target based on a literature search (**Table 1a**).

**Table 1a.** Putative targets of CBD in the CNS.

| Human Gene | Name | Effect | Mouse Ortholog | Cis eQTL (Tissue<sup>allele</sup>) | Ref |
| --- | --- | --- | --- | --- | --- |
| CACNA1G | Cav3.1 T-type calcium channel | Non-selective Antagonist | *Cacna1g* | NS | (69) |
| CACNA1H | Cav3.2 T-type calcium channel | Non-selective Antagonist | *Cacna1h* | Ctx<sup>B</sup>, Str<sup>B</sup> | (69) |
| CHRNA7 | Cholinergic receptor nicotinic alpha 7 subunit | Negative Allosteric Modulator | *Chrna7* | NS | (69) |
| DRD2 | Dopamine receptor D2 | Partial Agonist | *Drd2* | NS | (71) |
| ENT1 | Equilibrative nucleoside transporter 1 | Competitive Inhibition | *Ent1* | NS | (71) |
| HTR1A | Serotonin 1A receptor | Agonist | *Htr1a* | NS | (69, 71) |
| HTR2A | Serotonin 2A receptor | Partial Agonist | *Htr2a* | NS | (69, 71) |
| HTR3A | Serotonin 3A receptor subunit | Negative Allosteric Modulator | *Htr3a* | NS | (104) |
| GABRA1 | Gamma-aminobutyric acid type A receptor alpha 1 subunit ($\alpha 1\beta 2\gamma 2$) | Positive Allosteric Modulator | *Gabra1* | Ctx<sup>B</sup>, Hip<sup>B</sup>, Str<sup>B</sup> | (71) |
| GABRA2 | Gamma-aminobutyric acid type A receptor alpha 2 subunit ($\alpha 2\beta 3\gamma 2$) | Positive Allosteric Modulator | *Gabra2* | Ctx<sup>D</sup>, Hip<sup>D</sup>, Hyp<sup>D</sup>, Str<sup>D</sup> | (71) |
| GABRA4 | Gamma-aminobutyric acid type A receptor alpha 4 subunit ($\alpha 4\beta 2\gamma 2$) | Positive Allosteric Modulator | *Gabra4* | NS | (71) |
| GABRA5 | Gamma-aminobutyric acid type A receptor alpha 5 subunit ($\alpha 5\beta 2\gamma 2$) | Positive Allosteric Modulator | *Gabra5* | NS | (71) |
| GABRA6 | Gamma-aminobutyric acid type A receptor alpha 5 subunit ($\alpha 6\beta 2\gamma 2$) | Positive Allosteric Modulator | *Gabra6* | NS | (71) |
| GPR55 | G protein-coupled receptor 55 | Antagonist | *Gpr55* | NS | (69, 71) |
| KCNB1 | Potassium voltage-gated channel subfamily B member 1 (Kv2.1) | Antagonist | *Kcnb1* | Hyp<sup>B</sup> | (71) |
| KCNQ1 | Potassium voltage-gated channel subfamily Q member 1 (Kv7.1) | Antagonist | *Kcnq1* | NS | (71, 105) |
| KCNQ2 | Potassium voltage-gated channel subfamily Q member 2 (Kv7.2) | Positive Allosteric Modulator | *Kcnq2* | Ctx<sup>B</sup>, Hip<sup>B</sup>, Hyp<sup>B</sup>, Str<sup>B</sup> | (71, 105) |
| KCNQ3 | Potassium voltage-gated channel subfamily Q member 3 (Kv7.3) | Positive Allosteric Modulator | *Kcnq3* | NS | (71, 105) |
| KCNQ4 | Potassium voltage-gated channel subfamily Q member 4 (Kv7.4) | Positive Allosteric Modulator | *Kcnq4* | NS | (71, 105) |
| KCNQ5 | Potassium voltage-gated channel subfamily Q member 5 (Kv7.5) | Positive Allosteric Modulator | *Kcnq5* | Ctx<sup>B</sup>, Hip<sup>B</sup>, Hyp<sup>D</sup>, Str<sup>B</sup> | (71, 105) |
| PPARG | Peroxisome Proliferator Activated Receptor Gamma | Agonist | *Pparg* | NS | (69, 71) |
| SCN1A | Sodium voltage-gated channel alpha subunit 1 (Nav1.1) | Non-selective Antagonist | *Scna1* | NS | (71) |
| SCN2A | Sodium voltage-gated channel alpha subunit 2 (Nav1.2) | Non-selective Antagonist | *Scn2a* | Hip<sup>D</sup>, Str<sup>D</sup> | (71) |
| SCN3A | Sodium voltage-gated channel alpha subunit 3 (Nav1.3) | Non-selective Antagonist | *Scn3a* | Ctx<sup>B</sup>, Str<sup>B</sup> | (71) |
| SCN8A | Sodium voltage-gated channel alpha subunit 8 (Nav1.6) | Non-selective Antagonist | *Scn8a* | NS | (71) |
| SCN9A | Sodium voltage-gated channel alpha subunit 9 (Nav1.7) | Non-selective Antagonist | *Scn9a* | Ctx<sup>B</sup>, Hyp<sup>B</sup>, | (71) |
| TRPA1 | Transient receptor potential ankyrin type 1 | Agonist | *Trpa1* | NS | (69) |
| TRPM8 | Transient receptor potential subfamily M | Antagonist | *Trpm8* | NS | (69) |
| TRPV1 | Transient receptor potential vanilloid 1 | Agonist | *Trpv1* | NS | (69, 71) |
| VDAC1 | Voltage-dependent anion channel 1 | Direct Channel Blocker | *Vdac1* | NS | (69) |

For metabolic enzymes we limited our targets to those directly involved in THC and CBD metabolism (*e.g.*, Cytochrome P450 genes CYP2C9, CYP2C19 and CYP3A; fatty acid binding proteins or FABPs) (72-74). There is no single best ortholog for human P450 genes in mice as they have a greater number of P450 genes due to expansions within gene clusters. This is especially true for the *Cyp2c* cluster. Although *Cyp3a11* is often thought to be the primary functional mouse analog of human CYP3A genes, *Cyp3a13* and *Cyp3a25* can also contribute to liver metabolism. For these reasons we evaluated all liver-expressed *Cyp2c* and *Cyp3a* paralogs in BXD mouse liver datasets (**Table 1b**).

Targets were screened for possible functional expression differences between B6 and D2 strains using GeneNetwork (75). GeneNetwork (genenetwork.org) is an open online platform containing analysis tools and datasets, including genotype and brain gene expression data for BXD recombinant inbred (RI) mice. BXDRI are derived by intercrossing and inbreeding B6 and D2 mice (76). Quantitative trait loci (QTL) mapping of transcript or protein expression data in the BXDs can be used to identify genes whose expression is regulated by genetic variants segregating between B6 and D2. These genes are good candidates for causing and/or mediating innate strain differences in related behavioral responses as the expression differences in these genes are directly caused by genetic variation as opposed to environmental factors, consequences of drug exposure, or downstream actions of other genes.

We used the *Get Any:* search function to retrieve peak QTL locations in BXD hippocampus [*Hippocampus Consortium M430v2 (Jun06) RMA*], cortex [*HQF BXD Neocortex ILM6v1.1 (Dec10v2) RankInv*], hypothalamus [*INIA Hypothalamus Affy MoGene 1.0 ST (Nov10)*, *INIA Hypothalamus Affy MoGene 1.0 ST (Nov10) Female*, *INIA Hypothalamus Affy MoGene 1.0 ST (Nov10) Male*], striatum [*HQF Striatum Affy Mouse Exon 1.0ST Gene Level (Dec09) RMA*, *HQF BXD Striatum ILM6.1 (Dec10v2) RankInv)*], liver [Liver mRNA & Dataset = UTHSC BXD Liver RNA-Seq Avg (Oct19) TPM Log2, EPFL/LISP BXD CD Liver Affy Mouse Gene 1.0 ST (Aug18) RMA] data sets for all targets in **Table 1** using gene symbols for mouse homologs. An additional search for protein QTLs (pQTLs) was also performed for metabolic genes (*e.g.*, fatty acid binding proteins and cytochrome P450 genes) using a BXD control liver proteome database [Liver Proteome & Dataset = EPFL/ETHZ BXD Liver Proteome CD (Nov19)]. To ascertain whether gene or protein expression of each target was regulated by variants located within or near each target gene—so called local or cis expression quantitative trait loci or cis eQTLs—we retained genes with peak QTLs located within 2 Mb of the gene locus. A QTL was considered worthy of further attention when the associated LOD (logarithm of the odds) score was greater than 2 (*i.e.*, probability of the QTL is 100 times more probably than its absence).

## RESULTS

### CBD can attenuate or enhance THC-induced reductions in motor activity dependent on dose, time, sex, and strain

Motor response was quantifed as the time spent mobile in the open field at 0, 30, and 75 min following treatment. Initial and repeated sensitivity were calculated for B6 and D2 females and males for each treatment (*i.e.*, VEH, THC, or THC + CBD mixtures) and time point. For each group, we first determined whether there was a significant difference between any treatment upon initial or repeated exposure. Significant treatment effects were further evaluated within each group using Dunnetts’s post-hoc tests to determine whether any treatments influenced motor activity relative to VEH and whether any THC + CBD combinations influenced motor activity relative to THC alone.

#### CBD dose modulates the initial sensitivity to the hypolocomotor effects of THC in D2 females only

Comparison of treatment group means by ANOVA revealed a significant effect of treatment (*p* < 0.01) on acute motor response for all groups at 30 and 75 min following exposure (**Supplemental Table 2**). Immediately following injection (i.e., 0 min), a significant effect (*p* < 0.05) of treatment on motor response was only evident in D2 females (**Supplemental Table 2**).

Relative to VEH, all treatments exhibited significant inhibition of motor activity at 30 min (*p* < 0.001) and 75 min (*p* < 0.001) post-injection (**Figure 2A**) in B6 mice of both sexes. Similarly, all treatments showed significant inhibition of motor activity relative to VEH (**Figure 2C**) in D2 mice of both sexes at 30 min (p < 0.05) and 75 min (p < 0.05) following exposure. In addition, THC treatment caused a significant reduction in motor activity at time 0 relative to VEH (*p* < 0.05) in D2 females.

**Figure 2.**
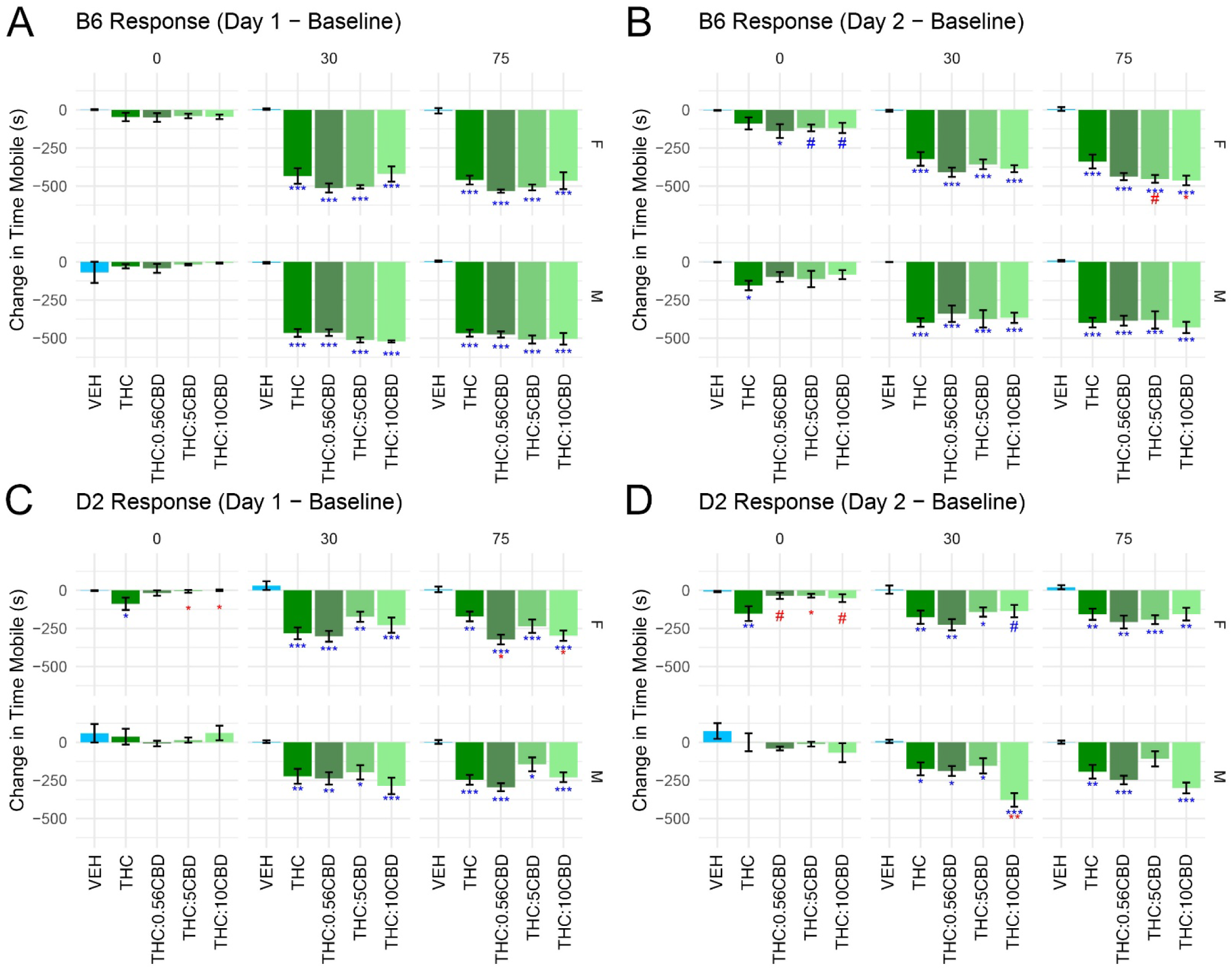
Acute motor response and rapid tolerance to treatment. Higher doses of CBD influence THC-induced motor responses in a strain and sex dependent manner. **(A)** Initial sensitivity in B6 mice. THC and THC + CBD mixtures produces significant inhibition of motor activity at 30- and 75 min post injection in both sexes compared to VEH. There is no additional effect of CBD on time mobile in the open field in B6 females or males when compared to THC alone. **(B)** Rapid tolerance in B6 mice. In females THC + CBD mixtures increase inhibition of motor activity at time 0 compared to VEH. Also in females, all treatment groups produce a significant reduction in time mobile at the 30- and 75 min time points when compared to VEH. In males, THC produces a significant reduction in time mobile at time 0 relative to VEH. In males, all treatment groups produce a significant reduction in motor activity at 30- and 75 min post-injection when compared to VEH. In females, higher doses of CBD (e.g. 5 and 10 mg/kg) enhance the inhibition of motor activity at 75 min post-injection compared to THC alone. There is no additional effect of CBD on the rapid tolerance to the inhibitory effects of THC on motor response in B6 males. **(C)** Initial sensitivity in D2 mice. When compared to VEH, THC and THC + CBD mixtures produce significant inhibition of motor activity at 30- and 75 min post injection in both sexes. In female D2 mice a significant reduction in time mobile compared to the VEH group is also observed at time 0, immediately following injection of THC. In female D2 mice, higher doses of CBD (*e.g.*, 5 and 10 mg/kg) reversed the THC-induced reduction in time mobile at time 0 and low and high amounts of CBD enhanced the THC-induced reduction in time mobile at 75 min post-injection. Relative to THC, there is no additional effect of CBD on time mobile in the open field in D2 males. **(D)** Rapid tolerance in D2 mice. In female D2 mice, THC produces a decrease in motor activity at time 0 and all treatments produce a decrease in activity at 30- and 75 min post-injection when compared to VEH. In male D2 mice relative to VEH, all treatments produced a significant decrease in motor activity at 30- and 75 min post-injection with the exception of THC.CBD5 at 75 min post-injection. When compared to THC, CBD reverses the inhibitory effects of THC at time 0 but has no additional effects at later time points in female D2 mice. In male D2 mice, CBD enhances THC-induced reductions in time mobile in the high CBD group at 30 min post-injection relative to THC alone. Significance thresholds: 0.15 < *p* > 0.05 = # (trend), *p* < 0.05 = *, *p* < 0.01 = **, *p* < 0.001 = ***. Significance indicated for Dunnett’s test with VEH as the control in blue and with THC as the control in red.

Importantly, THC + CBD combinations did not alter motor activity compared to THC in B6 mice of both sexes (**Figure 2A**). There was also no effect of CBD ratio on motor activity relative to THC treatment in D2 males. In D2 females, higher ratios of CBD (*e.g*., THC:5CBD and THC:10CBD) significantly (*p* < 0.05) attenuated the reduction in motor activity associated with THC at the 0 min timepoint (*p* < 0.05, **Figure 2C**) while low and high doses of CBD (*e.g.*, THC:0.56CBD and THC:10CBD) enhanced THC-induced reductions in motor activity at later timepoints (*e.g.*, 75 min post-injection) (**Figure 2C**).

#### CBD modulates hypolocomotor effects of THC after repeated exposure in D2 mice of both sexes and in female B6 mice

Similar to the results for initial sensitivity, a trending or significant effect of treatment on motor activity was still evident based on ANOVA for all groups and timepoints following two consecutive daily treatments; the exception being D2 males immediately following treatment at 0 min post-injection (**Supplemental Table 2**).

Relative to the VEH group, all THC + CBD combinations at 0 min post-injection increased inhibition of motor activity (*p-values* < 0.05 or trending) in B6 females (**Figure 2B**). At later time points all treatments resulted in a significant decrease (*p* < 0.001) in motor activity (**Figure 2B**) relative to VEH in B6 females. In B6 males, repeated exposure to THC also produced a signficant (all p-values < 0.05) reduction in motor activity at all time points relative to VEH (**Figure 2B**). In D2 females, repeated exposure to THC significantly decreased (*p* < 0.01) motor activity at most time points relative to VEH (**Figure 2D**). Repeated exposure to all treatments except THC.CBD5 produced a significant decrease in motor activity at both 30 min (*p* < 0.05) and 75 min (all *p*-values < 0.05) in D2 males relative to VEH (**Figure 2D)**.

When considering whether CBD combined with THC can alter behavior compared to THC alone, we found that the highest CBD ratio (*i.e.*, THC:10CBD) at 75 min post-injection was significantly (*p* < 0.05) associated with greater inhibition of motor activity relative to THC in B6 females (**Figure 2B)**. At the same time point, THC:5CBD treatment also trended towards greater inhibition of motor activity in B6 females relative to THC alone (**Figure 2B)**. In contrast, there was no effect of CBD dose on motor activity relative to THC in B6 males **(Figure 2B**). In D2 females, at time 0 and relative to THC alone, all three combinations of THC and CBD appeared to attenuate the inhibitory effects of THC on motor activity (*p-values* < 0.05 or trending), however, a significant effect of CBD dose relative to THC was not detected at later time points following repeated injection in D2 females (**Figure 2D**). Repeated exposure to the highest dose of CBD (*i.e.*,THC:10CB) produced a significant (*p* < 0.01) reduction in motor activity at the 30 min timepoint relative to THC alone in D2 males (**Figure 2D**).

### Modulation of THC-induced hypothermia by CBD is primarily observed in D2 mice with the direction of modulation dependent on time, dose, and sex

Initial sensitivity and rapid tolerance to the hypothermic effects of THC alone or in combination with CBD were assessed by measuring changes in body temperature between a baseline VEH injection and a single or repeated injection of VEH, THC, or THC + CBD mixtures at 0, 15, 30, 60, and 120 minutes post-injection. Body temperature was measured at all timepoints using non-invasive RFID transponders implanted subcutaneously prior to treatment.

#### CBD dose modulates initial sensitivity to THC-induced hypothermia in D2 but not B6 mice

ANOVA revealed a trending or significant effect of treatment (*p* < 0.01) on hypothermia for most groups at 0, 15, 30, 60 and 120 min following an acute exposure (**Supplemental Table 2**). Exceptions occurred immediately following injection (i.e., 0 min), where no effect of treatment was detected for B6 males and D2 mice of both sexes (**Supplemental Table 2**).

When compared to VEH, significant effects of treatment on hypothermia were not detected until 15 min post-injection (**Figure 3A,C**). At 15 min, a trending or significant decrease in body temperature relative to VEH was detected for all three THC:CBD ratios in B6 females, THC:5CBD treatment in B6 males, THC treatment in D2 females, and all three THC:CBD ratios in D2 males. At the 30 and 60 min timepoints all treatments were significantly associated with a decrease in body tempature in both strains and sexes relative to VEH (**Figure 3A,C**). At the last timepoint (*e.g.*, 120 min post-injection) all treatments resulted in significant hypothermia relative to VEH in B6 mice of both sexes and in D2 females. However, only the highest ratios of THC:CBD (*e.g.*, THC:5CBD and THC:10CBD) remained trending or significantly associated with a decrease in body temperature relative to VEH at 120 min post-injection in D2 males.

**Figure 3.**
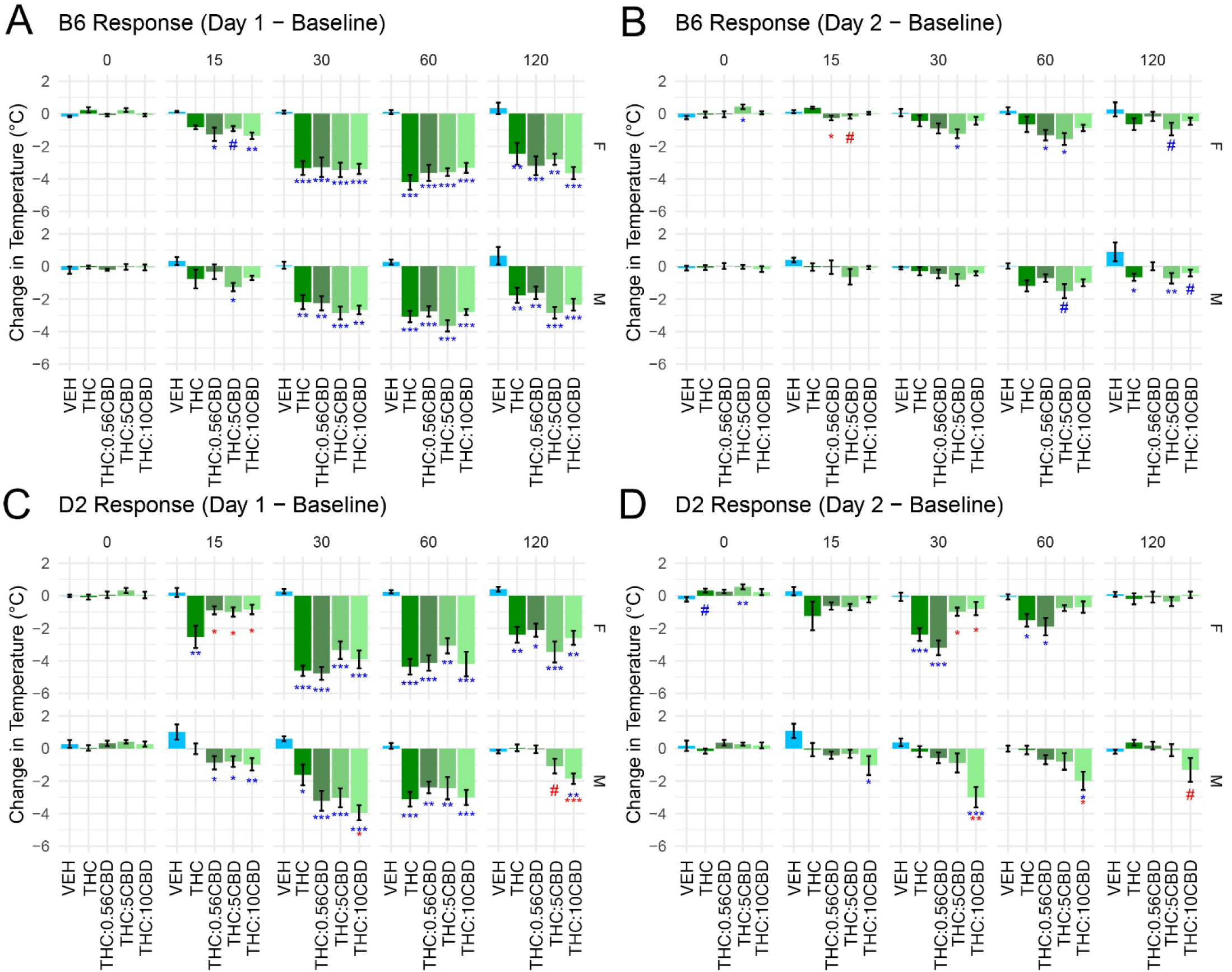
Acute hypothermic response and rapid tolerance to treatment. Higher doses of CBD influence THC-induced hypothermia in a strain and sex dependent manner. **(A)** Initial sensitivity in B6 mice. In B6 mice of both sexes, THC and THC + CBD mixtures are associated with a consistent and significant decrease body temperature starting at 30 min post-treatment relative to VEH. THC + CBD mixtures are also associated with a decrease in body temperature in females as early as 15 min following treatment relative to VEH. **(B)** Rapid tolerance in B6 mice. In both sexes, the effect of treatment groups on hypothermia relative to the VEH group is inconsistent following repeated exposure with only one or two treatment groups showing a significant decrease in temperature across time points. In female B6 mice 15 minutes following treatment, CBD appears to enhance hypothermia relative to THC alone at the low and middle dose, but not at the high dose. **(C)** Initial sensitivity in D2 mice. In D2 females, THC signifcantly decreases body temperature starting at 15 minutes post-treatment relative to VEH. All treatment groups produce a similar effect in D2 females at later time points relative to VEH. At 15 minutes following treatment, CBD significantly attenuates hypothethermia in D2 females realtive to THC. From 15 to 60 minutes following injection, all treatment groups are associated with a significant decrease in temperature in D2 males relative to VEH. In D2 males, the highest dose of CBD enhances hypothermia relative to THC alone at 30 min post injection. Also in D2 males and at the last time point, there is a trending or significant effect of CBD on hypothermia relative to THC, but only at the highest CBD doses. **(D)** Rapid tolerance in D2 mice. In D2 females THC and THC:0.56CBD still produce hypothermia relative to VEH following repeated exposure at 30 min and 60 min post-injection. At 30 min following treatment higher doses of CDB (e.g., THC:5CBD anf THC:10CBD) significantly attenuate hypothermia relative to THC. In D2 males, only the THC:10CBD treatment produces hypothermia at 30 and 60 min post-injection relative to VEH. Also in D2 males, the highest dose of CBD produces hypothermia at 30 min (*p* < 0.01), 60 min (*p* < 0.05), and 120 min (trend) relative to THC. Significance thresholds: 0.15 < *p* > 0.05 = # (trend), *p* < 0.05 = *, *p* < 0.01 = **, *p* < 0.001 = ***. Significance indicated for Dunnett’s test with VEH as the control in blue and with THC as the control in red.

Relative to THC, none of the THC:CBD ratios tested were associated with a change in body temperature in B6 females and males (**Figure 3A**). In contrast, all THC + CBD treatments significantly attenuated hypothermia at 15 minutes post-injection (*p* < 0.05) relative to THC alone in D2 females.

However, no significant effects of combined treatments on hypothermia were detected at later time points in D2 females **(Figure 3C)**. In contrast, trending or significant enhancement of THC-induced hypothermia was observed only at later time points following exposure D2 males **(Figure 3C)**. For example, the THC:10CBD treatment at 30 min was associated with enhanced hypothermia relative to THC alone as were the two highest THC:CBD ratios (*e.g.*, THC:5CBD and THC:10CBD) at the 120 min timepoint in D2 males (**Figure 3C**).

#### CBD dose modulates the repeated sensitivity to the hypolocomotor effects of THC in B6 females and D2 mice of both sexes

A trend or significant difference between treatment means following two consecutive days of treatment was detected by ANOVA for B6 females at all timepoints except at 120 min, for the 120 min timepoint only in B6 males, for the 0 min, 30 min, and 60 min timepoints in D2 females, and for all timepoints except the 0 min timepoint in D2 males.

In B6 females, when compared to VEH, body temperature was significantly increased by THC:5CBD treatment at 15 min and significantly decreased by both THC:5CBD treatment at 30 min and THC:0.56CBD and THC:5CBD treatment at 60 min (**Figure 3B**). There was a trend for decreased temperature relative to VEH in B6 females at the 120 min timepoint for the THC:5CBD treatment (**Figure 3B**). In B6 males a trend or significant decrease in temperature relative to VEH was detected at 60 min for the THC:5CBD treatment and for the THC, THC:5CBD, and THC:10CBD treatments at 120 min post-injection (**Figure 3B**). In contrast, following two consective daily treatments in D2 females, a trending or signficant decrease in temperature was detected both at 0 min for THC and THC:5CBD treatments and at 30 min for the THC and THC:0.56CBD treatments relative to VEH (**Figure 3D**). A significant decrease in temperature relative to VEH was only detected at the 15 min and 30 min timepoints in D2 males for the highest THC:CBD ratio (*e.g.*, THC:10CBD, **Figure 3D**).

No effect of CBD dose relative to THC treatment alone was detected for B6 males at any timepoint following repeated treatment. In B6 females, CBD dose was found to enhance hypothermia relative to THC, but only at 15 min after THC.0.56CBD and THC:5CBD treatments (**Figure 3B**). In contrast, CBD dose significantly attenuated THC-induced hypothermia at 30 min post-injection for the highest THC:CBD ratios (*e.g.*, THC:5CBD and THC:10CBD) in D2 females (**Figure 3D**). Paradoxically, THC:10CBD treatment in D2 males significantly or trended toward enhanced THC-induced hypothermia at 30, 60, and 120 mins following administration (**Figure 3D**).

### THC-induced antinociceptive responses are not modulated by CBD in B6 and D2 mice of either sex

Antinociception was quantified as tail flick latency in response to a thermal stimulus and was measured at 60 min following treatment. Initial and repeated sensitivity for tail flick latency were calculated for each sex, strain, and treatment and treatment effects were evaluated by ANOVA and Dunnett’s post-hoc tests as previously described.

#### CBD does not modulate initial sensitivity to THC-induced antinociception

Following an acute exposure, there was a significant effect of treatment on tail flick latency for all groups except D2 females (**Supplemental Table 2**). Following post hoc tests in B6 females and males, all treatment groups were found to be associated with a significant (*p* < 0.01) increase in tail flick latency relative to VEH **(Figure 4A)**. However, significant modulation of tail flick latency by combinations of THC and CBD relative to THC alone was not observed for B6 mice following acute treatment.

**Figure 4.**
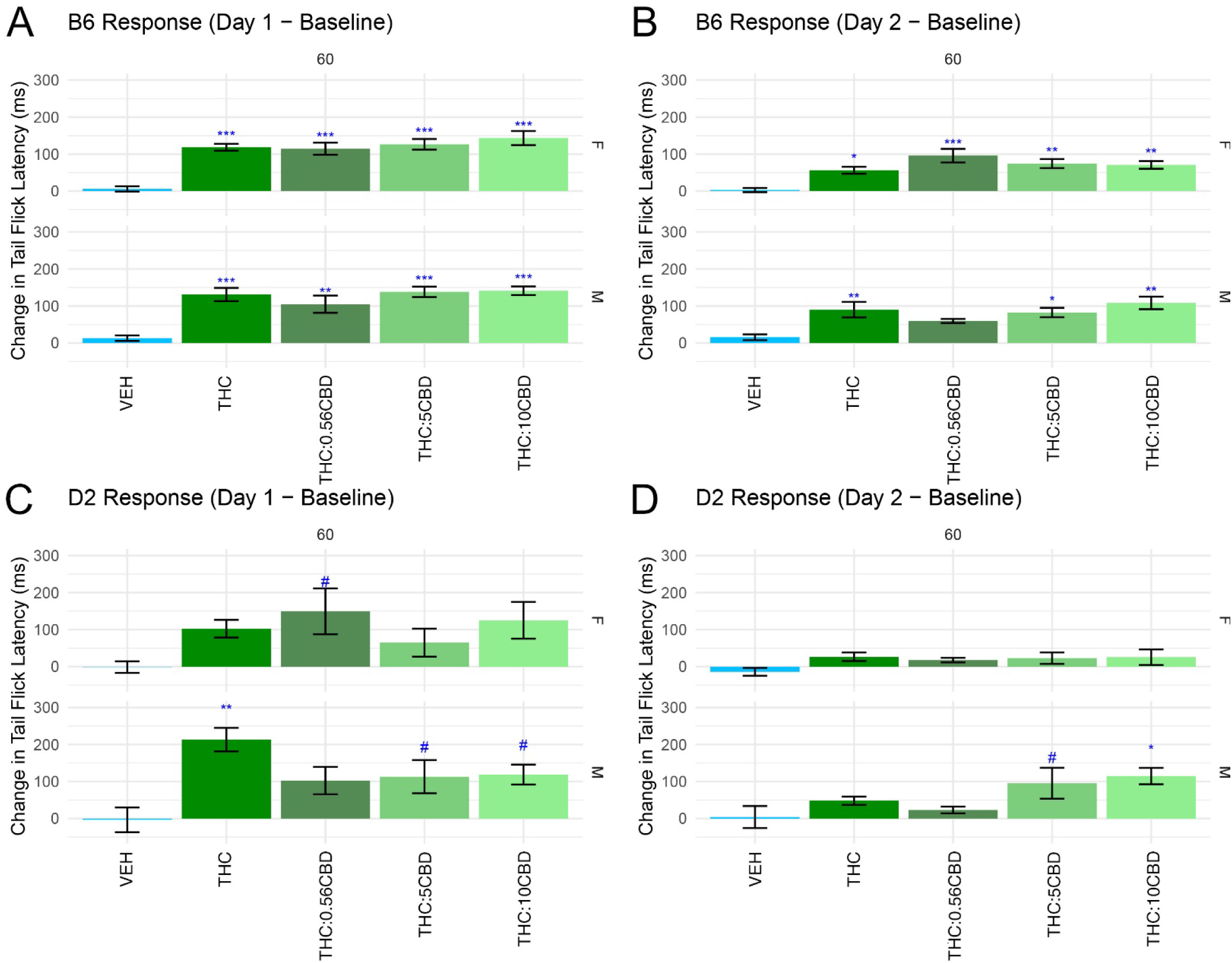
Acute antinociception and rapid tolerance to treatment. Although THC alone increases tail flick latency in response to a thermal stimulus in B6 mice of both sexes (**A**) and D2 males (**C**) upon first exposure, **t**here is no influence of CBD on THC-induced antinociception for initial sensitivity **(A)** or rapid tolerance **(B)** in B6 or D2 (**C**,**D**) mice. Upon repeated exposure, THC and most THC:CBD mixtures still result in a significant increase in tail flick latency relative to VEH in B6 mice of both sexes. However, this effect is not robustly observed in D2 mice with only the THC:5CBD and THC:10CBD treatments producing a trending or significant effect, respectively, relative to baseline in D2 males. Significance thresholds: 0.15 < *p* > 0.05 = # (trend), *p* < 0.05 = *, *p* < 0.01 = **, *p* < 0.001 = ***. Significance indicated for Dunnett’s test with VEH as the control in blue and with THC as the control in red.

In D2 males, treatment with THC alone or the highest ratio of CBD (*e.g.*, THC:5CBD and THC:10CBD) was associated with significant or trending increases in tail flick latency, respectively (**Figure 4C**). In D2 females, only the THC:0.56CBD treatment was associated with a trend towards increased tail flick latency following an acute exposure (**Figure 4C**).

#### No effect of CBD on repeated sensitivity to THC-induced antinociception

A significant effect of treatment on tail flick latency following repeated exposure was detected in all groups except for D2 females (**Supplemental Table 2**). Similar to the results following an acute exposure, all treatment groups except THC:0.56CBD in males were found to be significantly (*p* < 0.05) associated with an increase in tail flick latency relative to VEH following two consecutive exposures (**Figure 4B**) in B6 mice. In conrast, there was no difference in tail flick latency in D2 females for any treatment group relative to VEH following repeated exposure (**Figure 4D**). Only treatments with the highest ratio of CBD (*e.g*., THC:5CBD and THC:10CBD) produced a trending or significant increase in tail flick latency in D2 males relative to VEH, respectively (**Figure 4D**). Significant modulation of tail flick latency by THC + CBD combinations relative to THC alone was not observed for B6 or D2 mice following repeated exposure to treatment.

### Genetic analysis of candidate genes supports a pharmacodynamic mechanism of action for CBD modulation of THC-induced behavioral changes

The hypothermia, motor, and antinociception activity of THC is largely attributed to initial activation of CB1 receptors and subsequent secondary effects on signaling pathways in the CNS. The mechanism(s) of CBD are not as well defined with over 75 targets posited to mediate the biological activity of CBD, collated in a review by Marciniak et al. in 2025 (64). A list of plausible molecular targets of CBD was assembled based on pharmacokinetic studies of CBD in rodent models and the work of Bih et al. (69) and Manzoni et al. (71) followed by refinement based on follow-up searched of the current scientific literature. Probable endogenous CNS targets of CBD with *in vitro* activity (*e.g.*, EC50, IC50 or Ki) less than 10µM are shown in **Table 1A**. Probable metabolic enzyme targets of CBD directly involved in THC and CBD metabolism are shown in **Table 1B**. We used these targets as input to the GeneNetwork web resource (genenetwork.org, *see Methods*) to identify potential candidates that exhibit differential expression in the BXD recombinant inbred panel due to segregating variants between B6 and D2.

Of the plausible targets of CBD listed in **Table 1**, the expression patterns of 9 CNS genes—all of them ion channels, ligand-gated ion channels, or GPCRs—were significantly regulated by a cis eQTL in at least one brain tissue (**Table 1A**). The brain expression patterns of *Gabra2* (cortex, hippocampus, hypothalamus, and striatum) and *Scn2a* (hippocampus and striatum) were higher in BXDs that inherited the D2 allele of each gene relative to those with the B6 allele. In contrast, the expression patterns of *Cacna1h* (cortex and striatum), *Gabra1* (cortex, hippocampus, and striatum), *Kcnb1* (hypothalamus), *Kcnq2* (cortex, hippocampus, hypothalamus, and striatum), *Scn3a* (cortex and striatum), and *Scn9a* (cortex and hypothalamus) were higher in BXDs that inherited the B6 allele of each gene relative to those with the D2 allele. The allele effects on expression of *Kcnq5* were brain region dependent with higher expression associated with the B6 allele in cortex, hippocampus, and striatum and higher expression associated with the D2 allele in the hypothalamus.

Of the metabolic targets listed in **Table 1B**, only the cytochrome P450 genes exhibited genetic regulation of protein expression in liver. Of these, Cyp2c39, Cyp3a13, and Cyp3a25 demonstrated consistent differences at the transcript and protein expression level. Cyp3a13 had higher expression associated with inheritance of the B6 allele whereas Cyp2c39 and Cyp3a25 had higher expression associated with inheritance of the D2 allele. Overall, the parental allele effect within Cyp2c and Cyp3a gene clusters was inconsistent.

## DISCUSSION

### Modulation of THC-induced motor and hypothermia traits by CBD

Previously we showed differences in initial sensitivity to the acute behavioral effects (*e.g.,* motor activity, hypothermia, and antinociception) of a single high dose of THC (10 mg/kg) between strains and sexes (62). We largely replicated the results of the prior study. For example, B6 mice of both sexes are more sensitive to the inhibitory effects of THC on motor activity relative to D2 and females display greater sensitivity to the hypothermic effects of THC relative to males. Moreover, our current study demonstrates that co-treatment with CBD can modulate some of these THC-induced behavioral responses in a strain and sex dependent manner.

Specifically, our findings demonstrate that CBD significantly modulates the effects of THC on motor activity and hypothermia, but not antinociception, with distinct outcomes observed across different doses, times, and biological factors. Following the first exposure to treatment in female D2 mice, higher CBD levels (*e.g.*, 5 and 10 mg/kg) initially attenuated the ability of THC to reduce mobility in the open field within the first 10 min of injection and then paradoxically enhanced THC-induced motor suppression later at 75 min post-injection. A similar attenuation of THC-induced changes in motor activity immediately following treatment were also observed in D2 females for all CBD doses following repeated exposure. Enhancement of motor suppression by higher CBD levels (*e.g.*, 5 and 10 mg/kg) at later time points following treatment was also observed in B6 females, but only after repeated exposure. Conversely, only the highest CBD dose was able to amplify THC-induced suppression of motor activity at 30 min post-injection following repeated exposure in D2 males. There was no significant moderation by CBD in B6 males.

The hypothermic effects of THC were similarly modulated by CBD, showing distinct sex and strain dependent variations, highlighting the complexity of cannabinoid interactions. While there was not a significant effect of CBD in B6 mice, CBD consistently reversed THC-induced hypothermia across all doses 15 min following acute exposure in D2 females. In contrast, higher doses of CBD enhanced hypothermia at later time points, following acute or repeated exposure in D2 males indicating a dose dependent interaction on body temperature regulation.

### Potential underlying mechanism

CBD could modulate THC-induced changes in behavior through several mechanisms: 1) modulation of cannabinoid receptor activation by THC; 2) inhibition of THC metabolism; or 3) modulation of other specific endogenous targets. Given the CBD concentrations used in our study (*i.e.*, 0.56 to 10 mg/kg), we assumed an upper limit of 10 µM which falls below the range of possible allosteric regulation of CB1 and CB2 receptors. Therefore, we considered the evidence in support of inhibition of THC metabolism or specific action at sites other than cannabinoid receptors.

The effects of CBD on metabolism as well as THC-induced hypothermia, catalepsy, locomotion, and antinociception have been evaluated using different dosing paradigms and rodent models. For example, Jones and Pertwee (77) conducted the earliest study wherein adult male white mice (strain undefined) were pretreated for 30 min with CBD (50 mg/kg, *i.p.*) followed by THC (1.0 mg/kg, *i.v.*).

Pretreatment resulted in a 1.4 and 2-fold increase in 11-OH-THC (major metabolite) and THC, respectively, relative to THC alone. However, no change in catalepsy, defined as the immobility index, was measured within the limits of error in the assay (77). A pretreatment paradigm was also used by Bornheim *et al.* where male CF-1 mice were dosed with CBD (120 mg/kg, *i.p.*) and then administered THC (12 mg/kg, *i.v.* tail vein) two hours later resulting in a 26-fold and 14.5-fold increase of THC and 11-OH-THC metabolites in brain, respectively, relative to the THC controls (78). The interval between CBD (15-120 mg/kg, *i.p.*) pretreatment followed by THC (20 mg/kg or 12 mg/kg, *i.v.*) dosing was investigated by Reid and Bornheim (79) in male CF-1 mice. Immobility in the catalepsy test increased 40% when CBD (60 mg/kg, *i.p.*) was administered 60 minutes prior to 5 mg/kg THC. Brain concentrations of THC increased up to 3-fold in the 30-to-60-minute interval between CBD and THC administration. Of note, simultaneous administration of CBD and THC did not significantly increase THC levels in brain. Varvel and colleagues evaluated behavioral changes and THC levels in male ICR mice using CBD (3 or 30 mg/kg, *i.v.*) 10 minutes prior to administration of THC (0.3 or 3.0 mg/kg, *i.v.*) (80). The 3 mg/kg (*i.v.*) dose of CBD 10 minutes prior to administration of THC (0.1 to 3 mg/kg) did not affect behavior but at the 0.3 mg/kg threshold dose of THC, 30 mg/kg CBD significantly potentiated THC-induced antinociception but not spontaneous locomotion, catalepsy, or hypothermia. Only a CBD:THC ratio of 30:3 resulted in a significant increase in brain THC levels while blood THC levels were significantly increasing at the 30:3 and 30:0.3 ratios. Although the antinociceptive effects could be a result of increased THC levels the authors note that specific effects of CBD at other endogenous targets like fatty acid amide hydrolase (81), TRVP1 (82), or 5-HT_1A_ receptors (60) could also be involved.

Other studies explored the effects of simultaneous administration of CBD and THC on behavior. Hayakawa *et al*. (83) the effect of CBD (3-50 mg/kg, *i.p.*) and THC (1 mg/kg, *i.p.*) on locomotion and hypothermia in male ddY mice. A 3:1 CBD:THC ratio did not significantly affect locomotor activity or hypothermia. However, CBD:THC ratios of 10:1 and 50:1 significantly enhanced the hypothermic and cataleptic effects of THC. A different response pattern was observed by Todd *et al.,* (84) in male C57BL/6 mice wherein a 1:1 ratio of CBD:THC (10 mg/kg, *i.p.*) modestly reduced the hypothermic effects of THC alone at 30 minutes, but not at 90 or 360 minutes. In contrast, a modest enhancement in locomotor inhibition was measured 5 minutes post-injection but was insignificant thereafter (84). Another study by Kasten and colleagues (85) evaluated the effects of CBD:THC (20:10 mg/kg, *i.p.*) in combination versus THC (10 mg/kg, *i.p.*) alone as a function of age and sex in C57BL/6J mice. Locomotion in the THC and combined treatment groups was significantly reduced relative to control but there were no significant effects of age or sex, nor was there a significant difference between THC and combined treatment. Repeated exposures to THC and combined treatments abolished any differences between groups.

Taken together, the results of these experiments suggest that pretreatment with large doses (*i.e.*, ≥ 30 mg/kg) of CBD could lead to the inhibition of THC metabolism, although the consequences of this inhibition on behavior appear mixed. Interestingly, simultaneous co-administration of CBD and THC, which would normally occur during consumption of cannabis or derived products, appears not to have a profound impact on THC levels when compared to pretreatment with CBD. The impact of simultaneous CBD and THC administration appears to depend on dose, ratio, and genetic background. Moreover, these studies could not rule out the possibility of off-target effects of CBD on other endogenous targets making this the most likely mechanism underlying our observed differences in hypothermia and locomotion.

### Potential contribution of candidate genes to CBD modulation of THC-induced changes in hypothermia and locomotion

Our current study was designed to address two considerations: 1) simultaneous administration of THC:CBD; 2) constant THC level with variation of levels of CBD to explore dose response relationships; and 3) incorporation of different sexes and genetic backgrounds. We hypothesized that differences in response to THC associated with varying levels of CBD are caused by differences in sex and/or genetic background. To begin to understand the underlying molecular mechanisms of differential CBD modulation of THC-induced responses we assembled a list of plausible CNS and metabolic targets based on the current available literature and evaluated whether any candidates showed evidence of genetic regulation of gene expression in the BXD genetic population derived from B6 and D2 strains. These differentially expressed genes were used to propose possible mechanisms underlying the effects of CBD on THC-induced changes in motor and hypothermia traits.

We identified several CNS genes, notably calcium, sodium, and potassium voltage-gated ion channel subunit genes, and GABA receptor alpha subunits as possible targets of CBD that demonstrated differential expression based on inheritance of B6 or D2 alleles (**Table2a**). In addition to the CNS-associated genes, we also found some evidence of genetic regulation of key metabolic genes (**Table 2b**). This could be an indication that there are strain differences in the metabolic processing of both CBD and THC. However, the results are difficult to interpret. A major functional analog, *Cyp3a11*, did not show evidence of genetic regulation and the genetic regulation of transcript and protein expression in liver tissue was inconsistent within the *Cyp2c* and *Cypa3* clusters. There are few direct cytochrome P450 orthologs in mice and many paralogs do not have analogous metabolic function relative to their human counterparts. The potential contribution of metabolism cannot be entirely discounted; however, the aforementioned studies (83-85) using simultaneous THC:CBD injections suggest that this mechanism is unlikely in our experimental design.

**Table 2b.** Putative metabolic targets of CBD. No protein information was available in GeneNetwork for Cyp2c67, Cyp2c68, Cyp3a57, or Cyp3a59.

| Human Gene | Name | Mouse Ortholog or Cluster (QTL type/allele) | Ref |
| --- | --- | --- | --- |
| CYP2C9, CYP2C19 | cytochrome P450 2C9, 2C19 | *Cyp2c29*<sup>pQTL</sup>, *Cyp2c37*<sup>pQTL</sup>, *Cyp2c38*<sup>pQTL</sup>, *Cyp2c39*<sup>eQTL; pQTL</sup>, *Cyp2c40*<sup>pQTL</sup>, *Cyp2c50*<sup>pQTL</sup>, *Cyp2c54*<sup>pQTL</sup>, Cyp2c55, Cyp2c65, *Cyp2c66*<sup>eQTL</sup>, *Cyp2c67*<sup>eQTL</sup>, | (69) |
|  |  | *Cyp2c68*<sup>eQTL</sup>, *Cyp2c69*, *Cyp2c70*<sup>pQTL</sup> |  |
| CYP3A4,<br>CYP3A5 | cytochrome P450<br>3A4, 3A5 | *Cyp3a11*, *Cyp3a13*<sup>eQTL; pQTL</sup>,<br>*Cyp3a16*, *Cyp3a25*<sup>eQTL; pQTL</sup>,<br>*Cyp3a41a*<sup>pQTL</sup>, *Cyp3a41b*,<br>*Cyp3a44*<sup>pQTL</sup>, *Cyp3a57*<sup>eQTL</sup>,<br>*Cyp3a59* | (71) |
| FABP3 | Fatty acid binding<br>protein 3 | *Fabp3* | (71) |
| FABP5 | Fatty acid binding<br>protein 5 | *Fabp5* | (71) |
| FABP7 | Fatty acid binding<br>protein 7 | *Fabp7* | (71) |

The hypothermic response to THC is thought to be primarily mediated by the activation of CB1 receptors located on glutamatergic neurons in the hypothalamus (86) especially the preoptic anterior hypothalamus responsible for thermoregulation, (87, 88) and by CB1 receptors located on cortical glutamatergic neurons (45). Interestingly, CB1 rescue in glutamatergic neurons following genetic deletion did not fully restore hypothermic effects thus suggesting mechanisms in non-cortical glutamatergic and non-forebrain GABAergic circuits (86). Earlier studies using pharmacological inhibition indicated involvement of CB1 receptors on GABAergic neurons in the hypothermic response to THC (89) while also reporting synergistic effects of NMDA-CB1 receptor activation (90).

The inhibitory effect of THC on motor activity has been mapped to discreet regions of the CNS using conditional knockouts and CB1 restoration techniques in CB1 knockout mice (45, 86). Corticostriatal glutamatergic neurons have been implicated in regulating the hypolocomotor effects of THC while no significant effect was observed in forebrain CB1^-/-^GABAergic neurons. However, inhibition of locomotor activity was not completely abolished when cortical GABAergic interneurons expressing CB1 were present. The complexity of the signaling mechanisms involved locomotor regulation is highlighted by the work of Blázquez and colleagues (91). Using D1 receptor-CB1 knockout mice (Drd1-CB1-KO), the group identified GABAergic D1 medium spiny neurons as participants in the deficits associated with THC and control of motor coordination. The mapping of the effects of THC in various brain regions and cell types remains incomplete but it nevertheless provides possible site(s) for investigating how CBD can modulate the physiological responses to THC.

Identification of CBD targets that vary between B6 and D2 genetic backgrounds provides hypotheses for future investigation of the mechanism(s) whereby CBD could alter glutaminergic and GABAergic transmission and modulate THC-induced changes in hypothermia and motor activity. For example, the Cav_3.2_ T-type channels have been shown to control NMDA-sensitive glutamatergic receptor transmission (92). The K_v7.2_ potassium channels cluster with dopamine- and glutamate transporters and are involved in glutamatergic neurotransmission and excitotoxicity processes (93) and modulation of GABA release in hippocampal nerve terminals (94). The K_v7.5_ potassium channels control neuronal excitability through generation of the M-current, are activated by GABA (95) and are enriched in GABAergic neurons—including striatum medium spiny neurons (96) that are involved in motor control and inhibition of motor activity by THC. Sodium channels are also key players in regulating the neuronal excitation/inhibition balance that is critical for normal CNS function and thought to be interrupted by THC. In cortical structures and hippocampal granule cells Nav_1.2_ is highly localized in glutamatergic regions (97, 98). While Nav_1.3_ has lower levels of expression in brain, it is upregulated following neuronal injury (99).

While many of the targets of CBD in **Table 1a** show evidence of genetic regulation of expression and have the potential to modulate the behavioral effects of THC on hypothermia and hypolocomotion, *Gabra2* is the only target of CBD for which the causal variant underlying expression differences are known. A non-coding variant that is unique to the B6 strain and segregating in crosses like the BXDs where B6 is a parent, causes a ∼2-fold reduction in mRNA and protein levels (100) with a corresponding effect on behavior. For example, the variant in *Gabra2* has been shown to be associated with differential sensitivity to methamphetamine (101), differences in opioid withdrawal traits (102), and is a genetic modifier of Dravet syndrome in mice (*e.g.*, severe epilepsy associated with *SCNA1* haploinsufficiency in humans modeled as heterozygous deletion of *Scna1* in murine models (103). Given the abundance of GABA-A receptors containing the α2 subunit in brain, it is plausible that D2 mice that have a greater number of this receptor type relative to B6 and might therefore be more sensitive to CBD modulation. Additional studies using genetically engineered mice and/or pharmacological manipulation of GABA-A receptors would be needed to test this hypothesis.

### Study limitations

Our genetic analysis provides potential new pathways and experimental strategies that may help identify regulators of THC responses in the presence of CBD. However, this study has some important limitations. While we have enough subjects to detect a moderate to large effect of CBD on THC-induced behavioral changes, we cannot detect smaller but potentially meaningful cannabinoid interactions. Additionally, this study primarily focused on THC and CBD in limited combinations whereas cannabis and cannabis derived products consist of hundreds of different possibly bioactive compounds that may act in additive or synergist manner to influence behavior.

Future research should explore the broader chemical complexity of cannabis to determine how various cannabinoids and terpenes influence therapeutic and psychoactive outcomes. Expanding studies to include a wider range of cannabinoid ratios and formulations could provide a more comprehensive understanding of their effects and improve personalized treatment approaches. We also used a single route of administration in this study, and different routes of administration may produce different results. Finally, although we were able to propose putative targets and mechanisms given prior research and our observations, we did not validate true causal relationships and also cannot completely rule out the involvement of metabolic/pharmacokinetic mechanisms.

## Conclusions

The observed strain- and sex-dependent variability underscores the importance of personalized approaches in cannabinoid research and therapy. Understanding how genetic and biological factors influence cannabinoid interactions could inform tailored therapeutic strategies, particularly in conditions requiring modulation of motor activity, thermoregulation, and pain sensitivity. Future studies should explore the underlying molecular mechanisms driving these differences and assess the clinical relevance of varying CBD-to-THC ratios while reducing the unwanted psychoactive effects.

## ACKNOWLEDGEMENTS

This research was supported by NIDA grant R073239075 to MKM and BMM.

